# Effect of time delay on the synchronization of excitatory-inhibitory neural networks

**DOI:** 10.1101/2020.08.30.274662

**Authors:** Hwayeon Ryu, Sue Ann Campbell

**Author notes:** Corresponding author Email addresses (Hwayeon Ryu), (Sue Ann Campbell).

## Abstract

We study a model for a network of synaptically coupled, excitable neurons to identify the role of coupling delays in generating different network behaviors. The network consists of two distinct populations, each of which contains one excitatory-inhibitory neuron pair. The two pairs are coupled via delayed synaptic coupling between the excitatory neurons, while each inhibitory neuron is connected only to the corresponding excitatory neuron in the same population. We show that multiple equilibria can exist depending on the strength of the excitatory coupling between the populations. We conduct linear stability analysis of the equilibria and derive necessary conditions for delay-induced Hopf bifurcation. We show that these can induce two qualitatively different phase-locked behaviors, with the type of behavior determined by the sizes of the coupling delays. Numerical bifurcation analysis and simulations supplement and confirm our analytical results. Our work shows that the resting equilibrium point is unaffected by the coupling, thus the network exhibits bistability between a rest state and an oscillatory state. This may help understand how rhythms spontaneously arise neuronal networks.

## 1. Introduction

Neuronal networks in the brain involve two fundamental types of neurons: excitatory neurons tend to promote the firing of action potentials in other neurons, while inhibitory neurons do the opposite [19, 22, 33]. The presence of both types of cells is thought to be important for the formation of network oscillation patterns such as bursting and clustering [7, 23, 37, 48, 46, 60].

Different network architectures can be found in different brain areas. Global inhibition, where the excitatory cells are sparsely connected but receive a common inhibitory input due to highly connected inhibitory cells, has been implicated in the formation of bursting oscillations in the thalamus [48, 49, 50]. Networks where reciprocal excitatory and inhibitory connections dominate are thought to be important in rhythm generation in the hippocampus [37, 60]. However, for networks in the cortex, coupling between excitatory cells is considered to be important [1]. Large scale networks in the cortex typically have local excitatory-inhibitory circuits with long range excitatory connections [32, 34].

Whether excitatory or inhibitory, synaptic coupling represents the communication of electrical information from one neuron to another. Inherent in this process are time delays. Conduction delay results from the time it takes for the electrical information to travel along the axon of one neuron to the synapse with another neuron. Synaptic delay results from processes an chemical reactions that occur at the synapse itself. We refer to the total effect of these two delays as the coupling delay.

Numerical simulation studies of neural network models indicate that time delays can be important in the formation of network behavior [2, 26, 31, 37, 45, 55, 57]. To understand the mechanisms behind such behavior, mathematical analysis is needed. Several different approaches to this problem can be taken, depending on the context. Model networks of neurons that are intrinsically oscillatory can be studied using a phase model, phase resetting curve or a Poincaré map approach [16, 5, 6, 36, 13], but other approaches exist [43, 59]. Continuum models have been useful for studying delay induced wave-propagation in large scale networks [27, 28, 47]. Stability and bifurcation analysis has proved useful for studying of networks of excitable neurons [8, 9, 10, 11, 17, 52, 15, 18, 41, 54, 44, 59]. Relaxation oscillators, either inherently oscillatory or excitable, can be studied via geometric singular perturbation theory [14, 25, 4, 38, 39, 51]. However, with a few exceptions [4, 38, 39, 51] the work above focuses on networks where all the neurons are of a single type (either excitatory or inhibitory) or the coupling is diffusive or sigmoidal rather than synaptic.

In our previous study [51], we considered a network of excitable, relaxation oscillator neurons, previously studied in [48, 49], where two distinct populations, one excitatory and one inhibitory, are coupled with time-delayed synapses. The excitatory population is uncoupled, while the inhibitory population is tightly coupled without time delay. Based on a geometric singular perturbation analysis for relaxation oscillators, our results showed delays play a key role in producing synchronized network behaviors. The analysis helps to explain how coupling delays in either excitatory or inhibitory synapses contribute to generating synchronized rhythms. Despite the richness of these results, our work was restricted to the network of neurons modelled as relaxation oscillators. Also, further analysis on other types of network behaviors, such as clustered patterns, is needed to obtain a more complete understanding of how different population rhythms arise as a result of the interaction between coupling delays, intrinsic properties of each cell and network architecture.

To extend previous work while relaxing the model limitations mentioned above, we study a general network of excitable, non-relaxation oscillators. In this network, there are two distinct populations, each of which includes a pair of excitatory and inhibitory neurons. To relax our previous assumption where the excitatory cells were uncoupled, we couple them through time-delayed synapses while leaving inhibitory cells coupled only with their respective excitatory ones. This allows the interaction among excitatory cells, which may result in the emergence of different network behaviors such as clustering. Based on the extended model, our goal is to provide the existence and stability conditions of differing population behaviors in terms of intrinsic properties of cells, the size of coupling delays, and strengths.

Our paper is structured as follows: Section 2 presents the models for a single cell and for the coupled network which will be considered in our study. Section 3 describes our analysis results including linear stability analysis. Section 4 presents supplementary numerical simulations along with numerical bifurcation analysis results using XPPAUT [21] and DDE-BIFTOOL [20]. We conclude with a discussion in Section 5.

## 2. The Models

We first describe the model equations corresponding to an uncoupled, single cell. There are two types: one for inhibitory cells and one for excitatory cells. Then we introduce the synaptic coupling between the cells, coupling delays, and network architecture to be considered.

### 2.1. Single Cell Model

The generalized equations for this model are as follows:

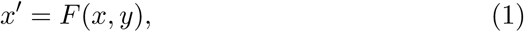

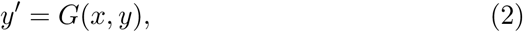

where 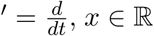, and *y* ∈ ℝ^*n*^. To simplify the analysis, we consider *n* = 1 in our study (see [49] for an example with *n* > 1). Also, we assume that the *x*-nullcline, *F* (*x, y*) = 0, is a cubic-*shaped* function, with left, middle, right branches, and *F* > 0 (*F* < 0) *below (above)* the *x*-nullcline curve. In addition, the *y*-nullcline is assumed to be a monotone increasing function *of x* that intersects with *x*-nullcline at the unique equilibrium point, and *G* > 0 (*G* < 0) *below (above)* the *y*-nullcline curve. Intersections of these curves determine equilibrium points of the system. See Figure 1.

**Figure 1:**
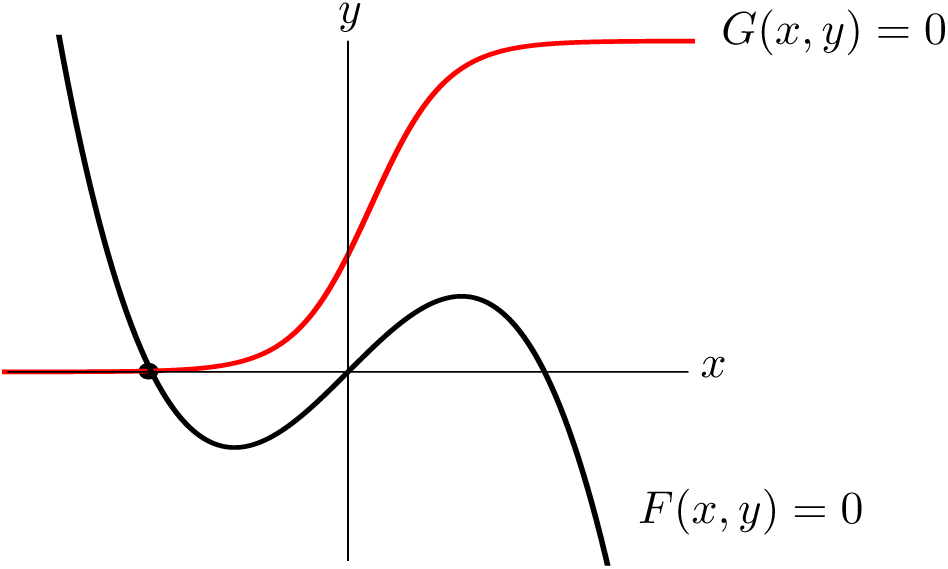
Nullclines for Eqs. (1)–(2). Black corresponds to the *x*-nullcline and red to the *y*-nullcline. The dot represents the unique equilibrium point for an excitable system.

Depending on the location of the equilibrium point (either left or middle branch) along the *x*-nullcline, the system is either *excitable* or *oscillatory*, respectively. For the excitable system, as can be seen in Figure 1, the equilibrium point is stable so no periodic solutions arise. In this situation, the equilibrium point is typically called the *resting equilibrium point* as the neurons are not active. However, if a sufficient amount of input, such as applied current, is applied to the excitable system, the equilibrium point can occur in the middle branch of the *x*-nullcline, resulting in an oscillatory system. A detailed discussion on oscillatory system behaviors with varying applied currents is provided in Appendix 6. Since many cells in the brain are active only when stimulated [19, 33, 22] our model cells are assumed to be excitable not oscillatory.

### 2.2. Synaptic coupling and network architecture

We consider networks with the architecture as shown in Figure 2. This is motivated by many brain regions, where there are local connections (within a region) between inhibitory and excitatory cells but global connections (between different region) are primarily between excitatory cells. This is the case, for example in the cortex, where inhibitory connections are primarily confined within a column, and even within a layer of a column, while the connections between columns or to different regions are excitatory [32, 34]. As a simple model of this, we consider a system with two excitatory and two inhibitory cells, where each *E*-*I* pair represents a different population. Thus there are no time delays in the connections between excitatory and inhibitory cells, but there are time delays in the connections between excitatory cells. We consider only two cells of each type to simplify the analysis. We will discuss how our work might be extended to larger networks in Section 5.

**Figure 2:**
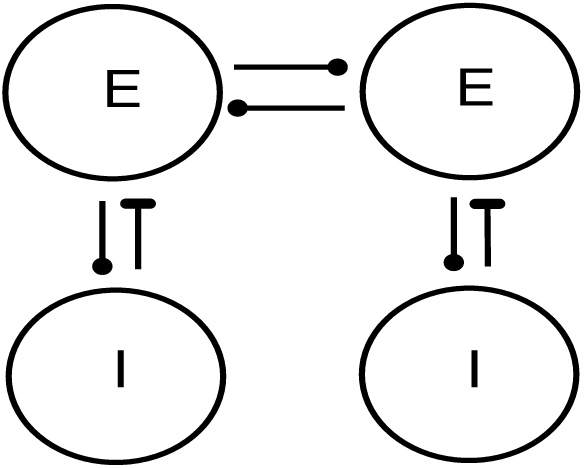
Schematic diagram of Excitatory-to-Excitatory network. Each *E*-cell interacts with each other, sending and receiving excitation, denoted by dot. Each *E*-cell also excites its coupled *I*-cell, which, in turn, inhibits its corresponding *E*-cell. Inhibitory connection is denoted by dash.

The equations corresponding to each *E*_*i*_ for *i* = 1, 2 in the network are

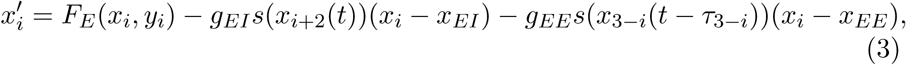

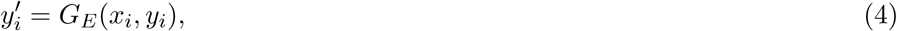

where *F*_*E*_, *G*_*E*_ correspond to *F, G* in Eqs. (1)–(2), and *g*_*EI*_ > 0 represents the maximal conductance of the *inhibitory* synapse, which can be viewed as coupling strength from the *I*-cell to its connected *E*-cell in the same population. The function *s* determines the synaptic coupling. It is a sigmoidal function which takes values in [0, 1]. Since the *I*-cell sends inhibition to the *E*-cells, *x*_*EI*_, the reversal potential for the synaptic connection, is set so that *x*_*i*_ − *x*_*EI*_ > 0, for the physiological range of values for *x*_*i*_. Thus we will assume *x*_*i*_ is less than the *x*-value of the resting equilbrium point for the uncoupled cell. We denote *x*_*inh*_ ≡ *x*_*EI*_. In a similar way, *g*_*EE*_ > 0 represents the maximal conductance of the *excitatory* synapse from the *E*-cell in the different population. *x*_*EE*_ represents the reversal potential for the excitatory connection, which is set so that *x*_*i*_ − *x*_*EE*_ < 0, for most of the physiological range of values for *x*_*i*_. Thus we will assume that *x*_*EE*_ is a small positive value. Finally, *τ*_3*−i*_ for *i* = 1, 2 denotes the delay in the excitatory synapse from *E*_3*−i*_ to *E*_*i*_ cells in different populations. Note that these are the only coupling delays in the network to be considered.

The model equations for *I*_*j−*2_ for *j* = 3, 4 are similarly given by

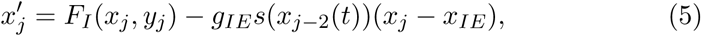

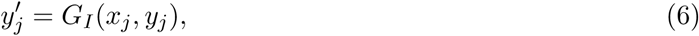

where *g*_*IE*_ denotes the maximal conductance of the excitatory synapse from *E* to *I* within the same population. For analysis simplicity, we assume *g*_*EI*_ = *g*_*IE*_ < *g*_*EE*_ but we could consider a different case for these strengths. The reversal potential for the excitatory synapse, denoted by *x*_*IE*_, is chosen so that *x*_*j*_ − *x*_*IE*_ < 0 for most of the physiological range of values for *x*_*j*_. We assume *x*_*exc*_ ≡ *x*_*EE*_ = *x*_*IE*_ but a different combination could be also considered for more complex network connections. It is assumed that there is no coupling delay in the synaptic connection from *E* to *I* within the same population.

Note that we do not incorporate chemical kinetics for synapses into our model. However, *τ*_1_ and *τ*_2_ include the effect of delays due to the chemical kinetics, as well as other factors. In addition, intrinsic properties of *E*- and *I*-cells, respectively, are identical in the system, in other words, *F*_*E*_ and *G*_*E*_ are the same for both *E*-cells, while *F*_*I*_ and *G*_*I*_ are the same for both *I*-cells.

Recall that an excitable cell stays at its stable equilibrium point if there is no an external synaptic input applied to the cell. The effect of this input depends on the type of coupling. For example, since *x*_*i*_ −*x*_*inh*_ > 0, inhibitory coupling decreases 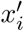 in (3) while excitatory coupling increases 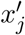 in (5), so long as *x*_*j*_ − *x*_*exc*_ < 0.

## 3. Model Analysis

### 3.1. Existence of equilibrium points

Let 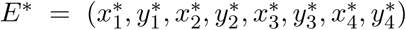 be an equilibrium point of the system (3)–(6). We assume that it is symmetric, i.e., in the form 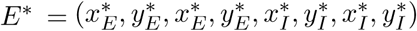. Plugging *E*^*∗*^ into the system yields that

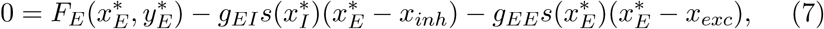

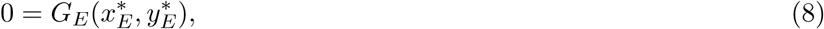

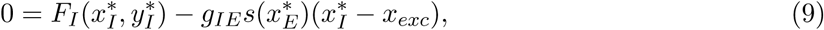

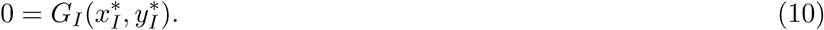

If there is no coupling, *g*_*EE*_ = *g*_*EI*_ = *g*_*IE*_ = 0, then there is an equilibrium point where each neuron is at its resting equilibrium point: 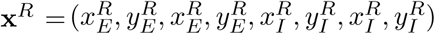 where 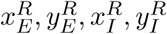 satisfy

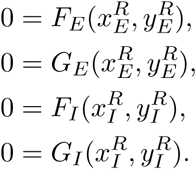

Typically the synapses are not active when the neuron is at rest, thus we will assume that 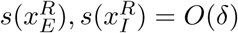 where 0 < *δ* «1. A simple perturbation argument then shows that when there is coupling in the network with *g*_*EE*_, *g*_*IE*_, *g*_*EI*_ = *O*(1) (or smaller) the resting equilibrium point persists and is given by **x**^*R*^ + *O*(*δ*).

Other equilibrium points may also exist. To see this, consider a simplified situation when there is no inhibition, that is, *g*_*EI*_ = 0. Then 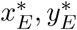 satisfy

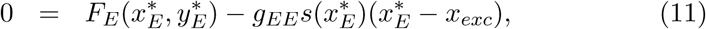

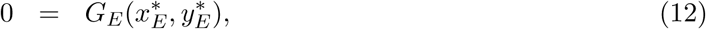

while the equations for 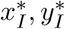 are unchanged, see (9)–(10). We can visualize the values of 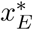 and 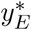 via intersections of the nullclines as shown in Fig. 3(a). The effect of the coupling term *I*_*EE*_ ≡ *g*_*EE*_*s*(*x*_*E*_)(*x*_*E*_ − *x*_*exc*_) can be understood as follows. Recall that *g*_*EE*_ > 0 and *s*(*x*_*E*_) ≥ 0 for all *x*_*E*_. Thus *I*_*EE*_ = 0 if *x*_*E*_ = *x*_*exc*_, negative if *x*_*E*_ < *x*_*exc*_ and positive otherwise. This can be seen in Fig. 3(a): the blue curve (*g*_*EE*_ > 0) intersects the black curve (*g*_*EE*_ = 0) at *x*_*exc*_, lies above it for *x* < *x*_*exc*_ and below it for *x* > *x*_*exc*_. However, if *x*_*E*_ is sufficiently small then *s*(*x*_*E*_) ≈ 0. Thus the black and blue curves are virtually identical for *x*_*E*_ sufficiently small. Since we assume 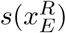 is small, the resting equilibrium point is almost unchanged by *g*_*EE*_, but if *g*_*EE*_ is large enough, other equilibrium points may exist as shown by the multiple intersection points of the blue and red curves in Fig. 3(a). The effect of the excitatory coupling on the inhibitory cell, 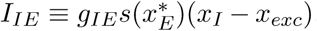, is similar but simpler. If 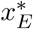 is small then 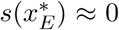 and 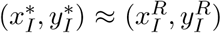. If 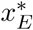 is large enough then *I*_*IE*_ will shift the *x*-nullcline up yielding moving 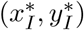 up and to the right. See Fig. 3(b).

**Figure 3:**
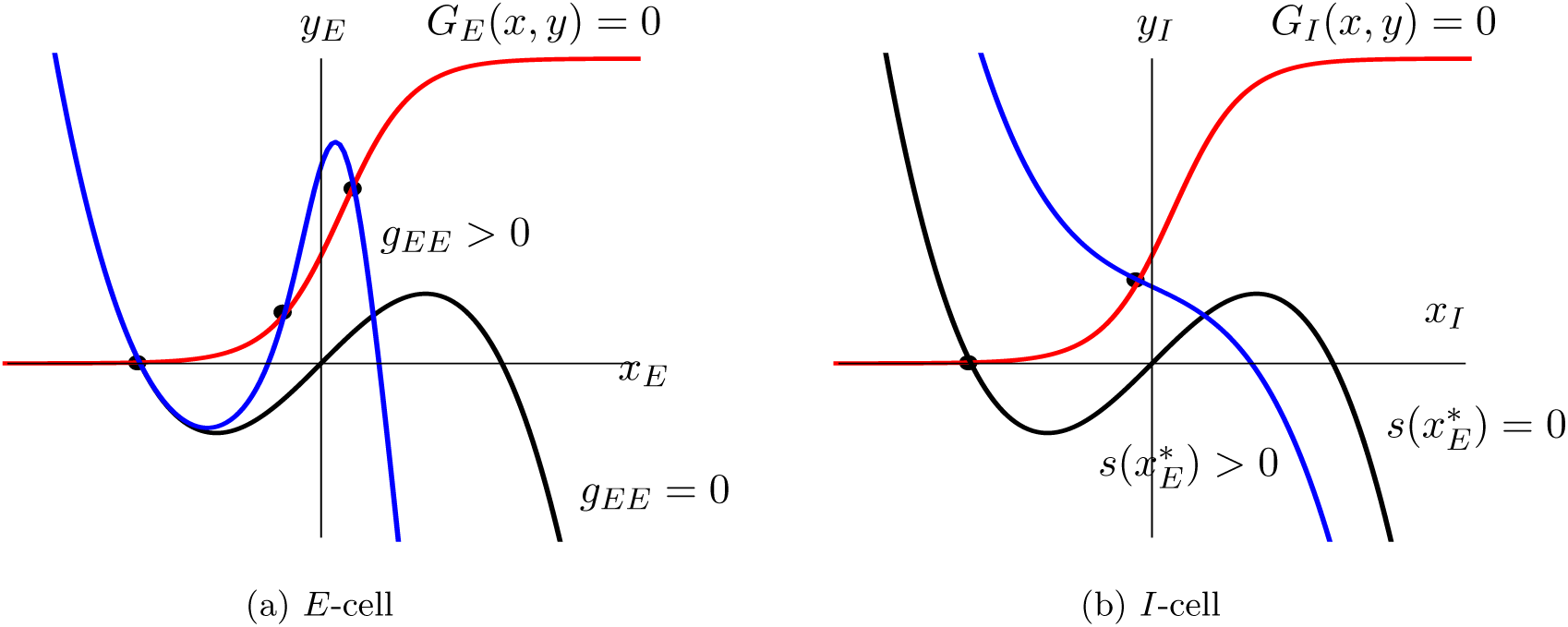
Effect of excitatory coupling on equilibrium points. Red curves correspond to *y*-nullclines. Black dots denote equilibrium points. (a) Black and blue curves correspond to *x*_*E*_ -nullcline (Eq. (11)) with *g*_*EE*_ = 0 and *g*_*EE*_ > 0, respectively. (b) Black and blue curves correspond to *x*_*I*_ -nullcline (Eq. (9)) with 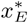 small enough that 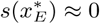 and large enough that 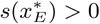, respectively.

If there is no coupling between the *E*-cells, *g*_*EE*_ = 0 but *g*_*EI*_, *g*_*IE*_ > 0, the only equilibrium point is the resting equilibrium point. This follows from the fact that for any *x*_*I*_, *C*_*EI*_ = *g*_*IE*_*s*(*x*_*I*_)(*x*_*E*_ − *x*_*inh*_) = 0 if *x*_*E*_ = *x*_*inh*_, is postive for *x*_*E*_ > *x*_*inh*_ and negative for *x*_*E*_ < *x*_*inh*_. Thus there can only be one equilibrium value for the *E*-cell and this satisfies 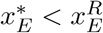. This means that 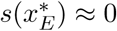, which in turn implies 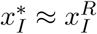.

If all couplings are present, the situation will be similar to the case with *g*_*EI*_ = 0. The only difference is that to obtain the extra equilibrium points, *g*_*EE*_ may need to be larger to overcome the inhibition.

### 3.2. Linear stability analysis

To conduct the linear stability analysis, we first write 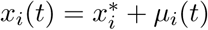 and 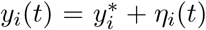 for *i* = 1, 2 and analogously, *x*_*j*_ (*t*), *y*_*j*_ (*t*) for *j* = 3, 4. Linearizing (3)–(6) about *E*^*∗*^ yields for *i* = 1, 2, *j* = 3, 4

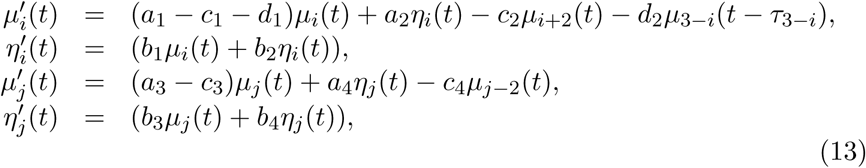

where 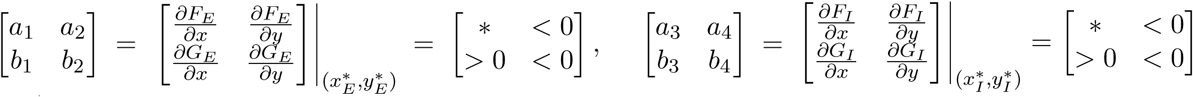

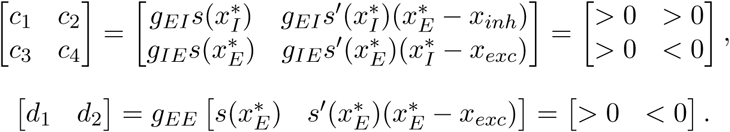

The signs for most coefficients are fixed as indicated based on our assumptions about the forms of the nonlinear functions. However, as can be seen from Figure 3, if *g*_*EE*_ > 0 the signs of *a*_1_ and *a*_3_ will depend on the value of 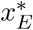 and 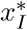. For the resting equilibrium point *a*_1_ < 0, *a*_3_ < 0 always. Further, 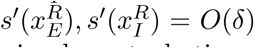 and *g*_*EE*_, *g*_*IE*_, *g*_*EI*_ = *O*(1) then *c*_*j*_, *d*_*j*_ = *O*(*δ*). Then, a simple perturbation argument shows that the stability of the resting equilibrium point is unaffected by the coupling.

The characteristic equation for this linear DDE is obtained by considering solutions of the form

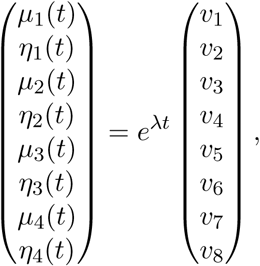

where *λ* = *β* + *iω* ∈ ℂ for *β, ω* ∈ ℝ. Such solutions will be nontrivial if and only if

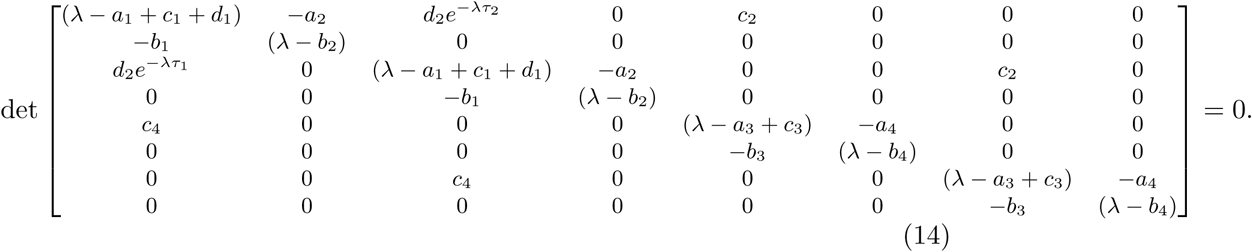

Expanding the determinant, we get the characteristic equation for the above 8-dimensional system:

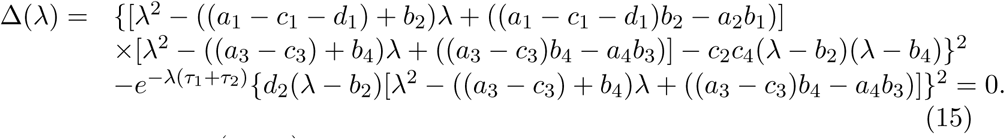

Let 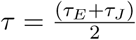 and *c* = *c*_2_*c*_4_ < 0. Then the characteristic equation (15) could be simplified as follows:

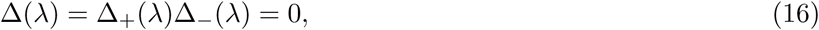

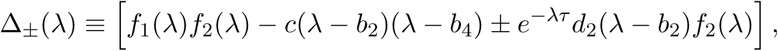

where *f*_1_(*λ*) = *λ*^2^ − (*a*_1_ − *c*_1_ − *d*_1_ + *b*_2_)*λ* + ((*a*_1_ − *c*_1_ − *d*_1_)*b*_2_ − *a*_2_*b*_1_) and *f*_2_(*λ*) = *λ*^2^ − (*a*_3_ − *c*_3_ + *b*_4_)*λ* + ((*a*_3_ − *c*_3_)*b*_4_ − *a*_4_*b*_3_). Since the coefficients of all terms in Δ_+_(*λ*) and Δ_+_(*λ*) are real, the roots of these functions come in complex conjugate pairs. Thus we may analyze the roots of the full characteristic equation by studying the roots of each factor separately.

#### 3.2.1. Single neuron case without any coupling: c_i_ = d_j_ = 0

We first consider the single neuron case where there is no coupling within the two *E*-*I* pairs as well as between two the *E* cells, that is, *g*_*EE*_ = *g*_*EI*_ = *g*_*IE*_ = 0. This implies *c*_*i*_ = *d*_*j*_ = 0 for *i* = 1, 2, 3, 4 and *j* = 1, 2 in (16). Thus we have

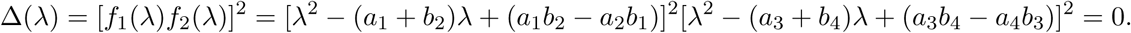

Note that the discriminant of the quadratic function, *f*_1_(*λ*), is (*a*_1_ − *b*_2_)^2^ + 4*a*_2_*b*_1_ whose sign depends on the specific values of the parameters. As discussed above, the only equilibrium point in this case is the resting equilibrium points, thus *a*_1_ < 0, *a*_3_ < 0. It follows that (*a*_1_ + *b*_2_) < 0 and (*a*_1_*b*_2_ − *a*_2_*b*_1_) > 0, which implies that if there are two real roots they are both negative. Even for complex roots, the first condition implies that the real part of complex conjugate roots will be negative. Similarly, the roots of *f*_2_(*λ*) all have negative real part. Thus, the equilibrium point *E*^*∗*^ is stable, which confirms that the neuron is unable to fire or oscillate without any synaptic coupling, representing an *excitable* system.

#### 3.2.2. Coupled system with a E-I coupling only: c_i_ ≠ 0, d_j_ = 0

Next we consider the coupled system but with a *E*-*I* coupling only to investigate how the stability of equilibrium point changes in response to the presence of inhibition from the *I* cells. This situation corresponds to *g*_*EE*_ = 0, *g*_*EI*_ > 0, *g*_*IE*_ > 0, thus we set *d*_*j*_ = 0 while keeping nonzero *c*_*i*_ in (16), which gives

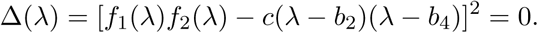

From the discussion in the previous section, the only equilibrium point in this case is the resting equilibrium point, thus *a*_1_ < 0, *a*_3_ < 0.

**I**. *λ* = 0 **case:** Plugging *λ* = 0 to the above equation results in

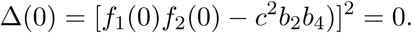

Since *f*_1_(0) = ((*a*_1_ − *c*_1_)*b*_2_ − *a*_2_*b*_1_) > 0, *f*_2_(0) = ((*a*_3_ − *c*_3_)*b*_4_ − *a*_4_*b*_3_) > 0 and *b*_2_*b*_4_ > 0, Δ(0) ≠ 0 for any *c* < 0. Thus, the characteristic equation cannot have a zero eigenvalue.

**II**. *λ* = ±*iω* **case:** Without loss of generality, we shall consider *λ* = *iω* only as the other case (*λ* = −*iω*) will be similar. Plugging *λ* = *iω* into (16) implies:

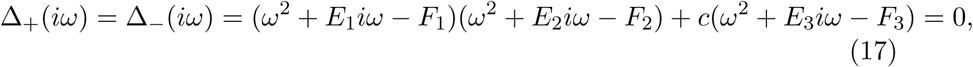

where *E*_1_ = (*a*_1_ − *c*_1_ − *d*_1_ + *b*_2_), *F*_1_ = ((*a*_1_ − *c*_1_ − *d*_1_)*b*_2_ − *a*_2_*b*_1_), *E*_2_ = (*a*_3_ − *c*_3_ + *b*_4_), *F*_2_ = ((*a*_3_ − *c*_3_)*b*_4_ − *a*_4_*b*_3_), *E*_3_ = (*b*_2_ + *b*_4_), *F*_3_ =^2^ *b*_2_*b*_4_. Given all the specified signs in (13) from Section 3.2, we know that *E*_*i*_ < 0, *F*_*i*_ > 0 for all *i* = 1, 2, 3.

Separating equation (17) into real and imaginary parts to rewrite in the form of Δ_*±*_(*iω*) = ℜ_*±*_(*iω*) + *i*𝔍_*±*_(*iω*) gives:

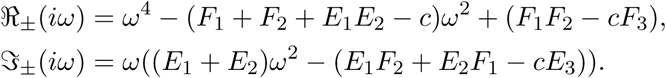

To satisfy Δ_*±*_(*iω*) = 0 for a nonzero *ω* ∈ ℝ ℝ, both ℜ ℜ_*±*_(*iω*) = 0 and 𝔍_*±*_(*iω*) = 0 should be satisfied. Solving for 𝔍_*±*_(*iω*) = 0 yields

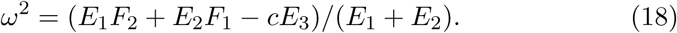

Since both the numerator and denominator in (18) are negative, such *ω*^2^ should exist. Also, because (*F*_1_ + *F*_2_ + *E*_1_*E*_2_ − *c*) > 0, (*F*_1_*F*_2_ − *cF*_3_) > 0 in ℜ_*±*_(*iω*), there are two positive roots for *ω*^2^. Using the quadratic formula, the two solutions could be obtained, however, none of which is the same as *ω*^2^ found in Eq. (18). This implies that there is no *ω*, for which ℜ_*±*_(*iω*) = 𝔍_*±*_(*iω*) = 0, i.e., there are no pure imaginary eigenvalues for (16). Thus, the equilibrium point *E* is asymptotically stable if no coupling between *E*-*E* is present despite the synaptic coupling between *E*-*I*. This result is consistent with our previous study [51], in which no oscillatory solution was found in the case of no delay in the *E*-*I* coupling.

#### 3.2.3. Coupled system with coupling delay: c_i_ ≠ 0, d_j_ ≠ 0, τ > 0

Now we study the impact of excitatory coupling on the solution behaviors by considering the fully coupled system with delay. Note that the corresponding characteristic equation with nonzero *d*_*j*_ and *τ* is given in (16).

**I**. *λ* = 0 **case:** To check the existence of a zero eigenvalue, we plug in *λ* = 0 in (16) to obtain

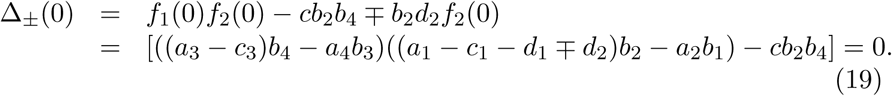

Ass ume that *a*_1_ < 0 and *a*_3_ < 0. Since (*a*_3_ − *c*_3_)*b*_4_ − *a*_4_*b*_3_ > 0, (*a*_1_ − *c*_1_ − *d*_1_ + *d*_2_)*b*_2_ − *a*_2_*b*_1_ > 0 and *c, b*_2_, *b*_4_ < 0, Δ_*−*_(0)0 since *d*_2_ < 0. However, there may exist *d*^*∗*^ ≡ *d*_2_ = [((*a*_3_ − *c*_3_)*b*_4_ − *a*_4_*b*_3_)((*a*_1_ − *c*_1_ − *d*_1_)*b*_2_ − *a*_2_*b*_1_) − *cb*_2_*b*_4_]*/*[*b*_2_((*a*_3_ − *c*_3_)*b*_4_ − *a*_4_*b*_3_)] < 0 such that Δ_+_(0) = 0, because the numerator is positive but the denominator is negative. Thus, there may be a zero eigenvalue, indicating that it is possible for the system to have multiple equilibrium solutions. This may also imply that the presence of excitatory synaptic coupling plays an important role in exhibiting other types of solutions in addition to the resting equilibrium solution.

Note that the existence of zero eigenvalue is valid even if there is no coupling delay (i.e., *τ* = 0) between *E*-*E* cells because *e*^*−λτ*^ = 1 for *λ* = 0 in (16). In addition, we could consider an even simpler case where the coupling between *E*-*I* is additionally removed by setting *c*_*i*_ = 0 for all *i* in Eq. (19). Then, the four-cell system reduces to a pair of coupled *E*-*E* cells only as two *I* cells are completely decoupled from this pair. Based on the same analysis, it follows that a zero eigenvalue can still exist with no coupling between the *E* and *I* cells. This result supports previous modeling studies [8, 9, 10, 11, 58], in which qualitatively different equilibrium solutions in neural networks were observed regardless of the presence of coupling delay.

**II**. *λ* = ±*iω* **case**: In addition, if we want to check whether the equilibrium loses its stability as the value of *τ* increases and also to find the critical delay value at which the delay-induced bifurcation occurs, let us consider the case of *λ* = ±*iω*.

First plugging *λ* = *iω* in (16), we get

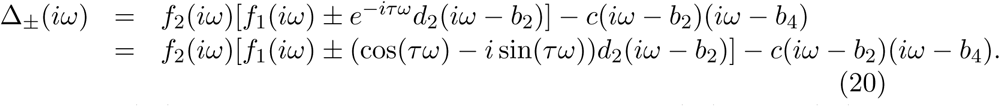

Sep arating (20) into real and imaginary parts, i.e., Δ_*±*_(*iω*) = ℜ_*±*_(*iω*) + *i*𝔍_*±*_(*iω*), and setting each part equal zero:

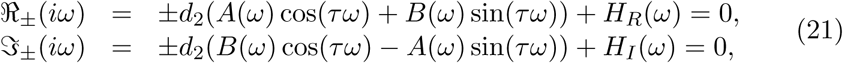

where

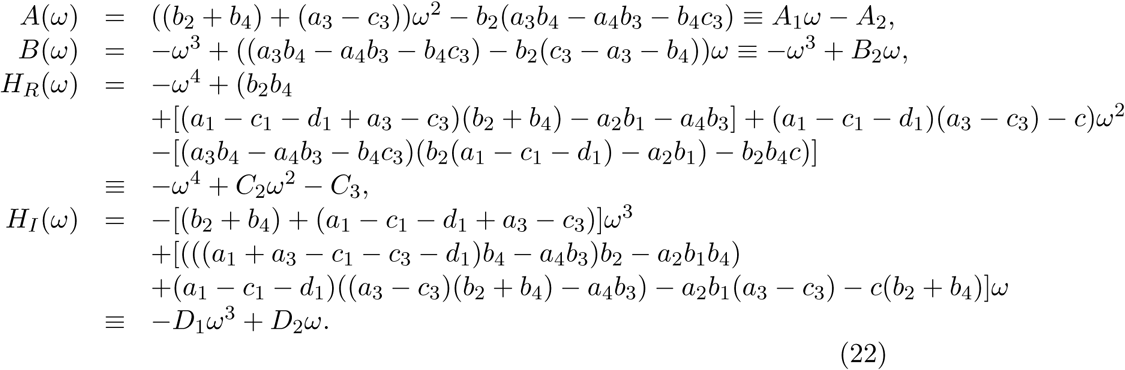

For Δ_+_(*iω*), we first gather all the terms of cos(*τω*) and sin(*τω*) by rearranging the above two equations in (21):

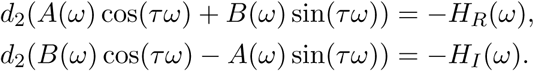

After algebraic manipulation, we can isolate cos(*τω*) and sin(*τω*) in terms of all other parameters.

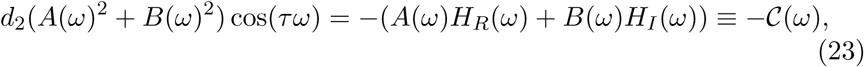

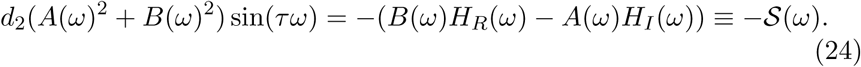

To eliminate *τ*, square the equations in (23)–(24), add and simplify to give

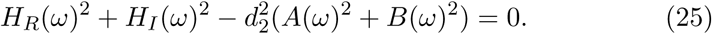

This is a degree-eight function in *ω* with coefficients that depend on all the parameters except *τ*. Dividing (24) by (23) results in tan(*τω*) = S(*ω*)*/*C(*ω*). However, this loses information about the signs of cos(*τω*) and sin(*τω*) that are in (23)–(24). Thus we introduce *y* = arctan(*u*) as the branch of the arctangent function with range 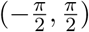. Note that this corresponds to cos(*y*) > 0 and that the function arctan(*u*) + *π* corresponds to cos(*y*) < 0. The other branches of the arctangent function are obtained from these two by adding multiple of 2*π*. As can be seen from (23), since *d*_2_ < 0 the sign of cos(*τω*) is determined by C(*ω*), and thus we define

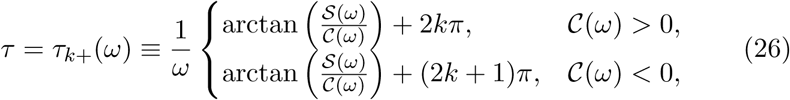

where *k* = 0, 1, ….

For Δ_*−*_(*iω*) the only difference is that *d*_2_ is replaced by −*d*_2_ in (23)-(24). Thus, in this case the signs of cos(*τω*) and C(*ω*) are the opposite. Hence we have

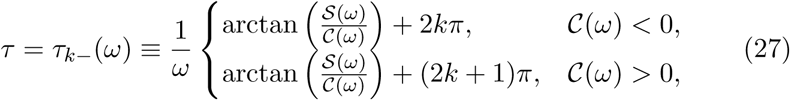

where *k* = 0, 1, Note that we do not take *k* < 0 as these branches yield *τ* < 0.

Let us finally consider a simpler case of no delay, as similarly discussed in zero eigenvalue section. One can do similar analysis and derive the same result in (25) by setting *τ* = 0 in (20)–(21) with the functions defined in (22). It can be easily shown that, for appropriate parameter values, there may exist pure imaginary eigenvalues for the characteristic equation associated with the coupled system with no delay.

#### 3.2.4. Eigenvector Analysis

To complete this section, we consider the form of the eigenvectors associated with the characteristic equation (16). First we rewrite the eigenvalue-eigenvector equation corresponding to the linearization (13) in a slightly different form, which corresponds to reordering the variables as (*µ*_1_, *η*_1_, *µ*_3_, *η*_3_, *µ*_2_, *η*_2_, *µ*_4_, *η*_4_).

Then solutions of (13) of the form **v***e*^*λt*^ satisfy

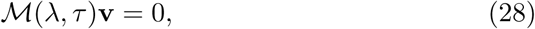

where

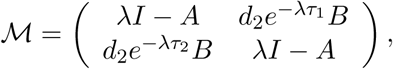

with *I* the 4 × 4 identity matrix and

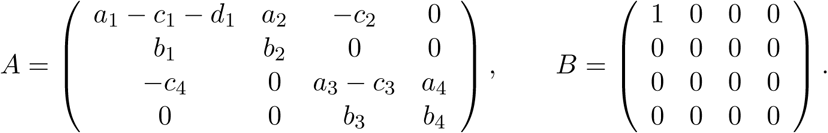

Nontrivial solutions will correspond to values of *λ* that satisfy the characteristic equation (16). Let *λ* be such a value and **v** the corresponding solution of Eq. (28). Define the 8 × 8 invertible matrix

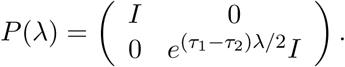

Then Eq. (28) implies

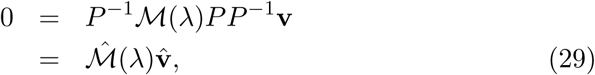

where 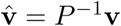,

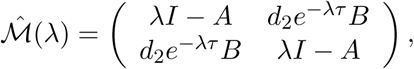

and *τ* = (*τ*_1_ +*τ*_2_)*/*2. The matrix 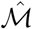 is exactly in the form considered by [59]. Following their approach, let *ξ* ∈ ℝ^4^. Then there is nontrivial vector

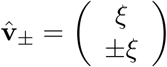

that satisfies Eq. (29) if and only if there is a nontrivial *ξ* that satisfies

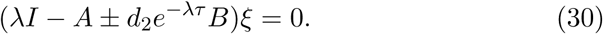

This latter is true if

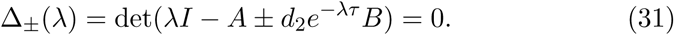

It follows that if *λ* is root of Δ_*±*_(*λ*) and *ξ* is a nontrivial vector satisfying (30) then the vector

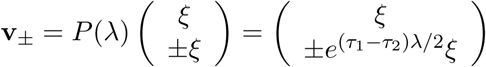

satisfies Eq. (28).

For example, suppose that *τ*_1_ = *τ*_2_. If *λ* = ±*iω* is a root of Δ_+_(*λ*) in (31) then the corresponding eigenvector will be (*ξ, ξ*)^*T*^, which will yield a periodic solution of (13) with *µ*_1_(*t*) = *µ*_2_(*t*), *µ*_3_(*t*) = *µ*_4_(*t*) and similarly for *ν*_*j*_. However, if *λ* = ±*iω* is a root of Δ_*−*_(*λ*) then the corresponding eigenvector will be (*ξ*, −*ξ*)^*T*^. The complex form of the solution will satisfy *e*^*i*(*ω*(*t*+*π/ω*))^**v** = −*e*^*i*(*ωt*)^**v**. Thus the corresponding periodic solution of (13) will satisfy *µ*_1_(*t* + *T/*2) = *µ*_2_(*t*), *µ*_3_(*t* + *T/*2) = *µ*_4_(*t*) where *T* = 2*π/ω*, and similarly for *η*_*j*_. So the *E*-cells are phase-locked with phase difference of *T/*2, as are the *I*-cells. A similar analysis if *τ*_1_*τ*_2_ shows that the corresponding solution will be phase locked with phase difference 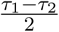, if *λ* = ±*iω* are roots of Δ_+_(*λ*), and with phase difference 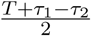 if *λ* = ±*iω* are roots of Δ_*−*_(*λ*).

## 4. Numerical Results

To further our study, we considered the following specific choice for the functions in (1)–(2):

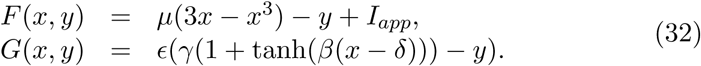

This model is inspired by that of [56]. It has a cubic nonlinearity as in for the FitzHugh-Nagumo model [24, 42], but with a nonlinearity in the equation for the “recovery” variable which is similar to that for a gating variable in a conductance-based model. We choose the same *F* and *G* for both excitatory and inhibitory cells.

The parameters were chosen as follows

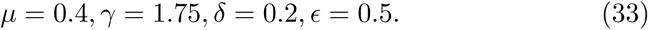

A bifurcation study of this model shows that with these parameters the parameter *β* could be used to switch the model from a class 2 (*β* = 1.5) to a class 1 (*β* = 2.0) oscillator, with *I*_*app*_ used as the parameter. Details can be found in Appendix. For both values of *β* the cell is excitable for *I*_*app*_ small enough, exhibits stable spiking behavior (has a stable limit cycle) for a middle range of *I*_*app*_ values and has a “high amplitude” stable equilibrium point for *I*_*app*_ large enough. Typical spiking solutions are show in the appendix, where it can be seen that that *x* varies in the range [−2, 1] for these solutions.

For our studies of coupled cells, we set *I*_*app*_ = 0 in all cells, so that all cells are excitable and the only input comes from the synapses with other cells. The synaptic coupling function is

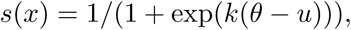

with *k* = 5 and *θ* = 0.1. The synaptic reversal potentials are *x*_*EE*_ = *x*_*IE*_ = 0.5, and *x*_*EI*_ = −2. With this choice *x*_*E*_ − *x*_*EI*_ > 0 for all *x* on a typical spiking solution while *xE/I* − 0.5 < 0 for most points of the typical spiking solution. Thus these choices give similar input to the cells as standard values in conductance based models. The coupling strengths were varied as shown below.

### 4.1. Hopf Bifurcations

To begin, we studied the case when there is no delay in the coupling. That is, we considered the system given by Eqs. (3)–(6), with *τ*_1_ = *τ*_2_ = 0, *F*_*E*_ = *F*_*I*_ = *F* and *G*_*E*_ = *G*_*I*_ = *G* where *F, G* are given by Eq. (32). We used XPPAUT [21] to carryout numerical bifurcation analysis of the model when the excitatory coupling strength is varied. As shown in Figure 4, there is one equilibrium point that appears to always exist and be asymptotically stable, at least for the range of *g*_*EE*_ we considered. This equilibrium point corresponds to all cells being at their resting equilibrium point. In other trials we considered *g*_*EE*_ ∈ [0, 700] and this equilibrium point still persisted. Note that, for our parameter values, 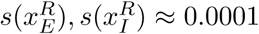, and 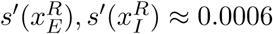. This confirms the analysis of Section 3.1: if the coupling strengths are *O*(1) with respect to the size of 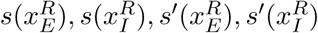 the resting equilibrium point exists and is asymptotically stable. At a critical value of the excitatory coupling, *g*_*EE*_, a pair of unstable equilibrium points of the form 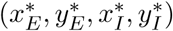, is created in a saddle-node (fold) bifurcation. For larger *g*_*EE*_, one of these equilibria is stabilized in a Hopf bifurcation. The other appears to always be unstable. The Hopf bifurcation is subcritical and gives rises to an unstable periodic orbit which is destroyed in a homoclinic bifurcation. Varying the other conductances *g*_*EI*_ and *g*_*IE*_ or the value of *β* did not affect the structure of the diagram, just the sizes of the equilibrium points and the region of existence of the periodic orbit. See Figure 5. In this figure we see a threshold for *g* = *g*_*EI*_ = *g*_*IE*_. If *g* < 1 the saddle node and Hopf bifurcation occur as the same value as when *g* = 0. Thus the connections between the *E* and *I* cells have little effect on the dynamics of the system. Above this threshold the inhibition of the *E* cells by the *I* cells has an effect: larger *g*_*EE*_ values are needed to cause the saddle node and Hopf bifurations. In these diagrams we kept *g*_*EI*_ = *g*_*IE*_, however, varying these independently showed the similar behavior. There are thresholds for these parameters such that both must be above their threshold for inhibition to have an effect.

**Figure 4:**
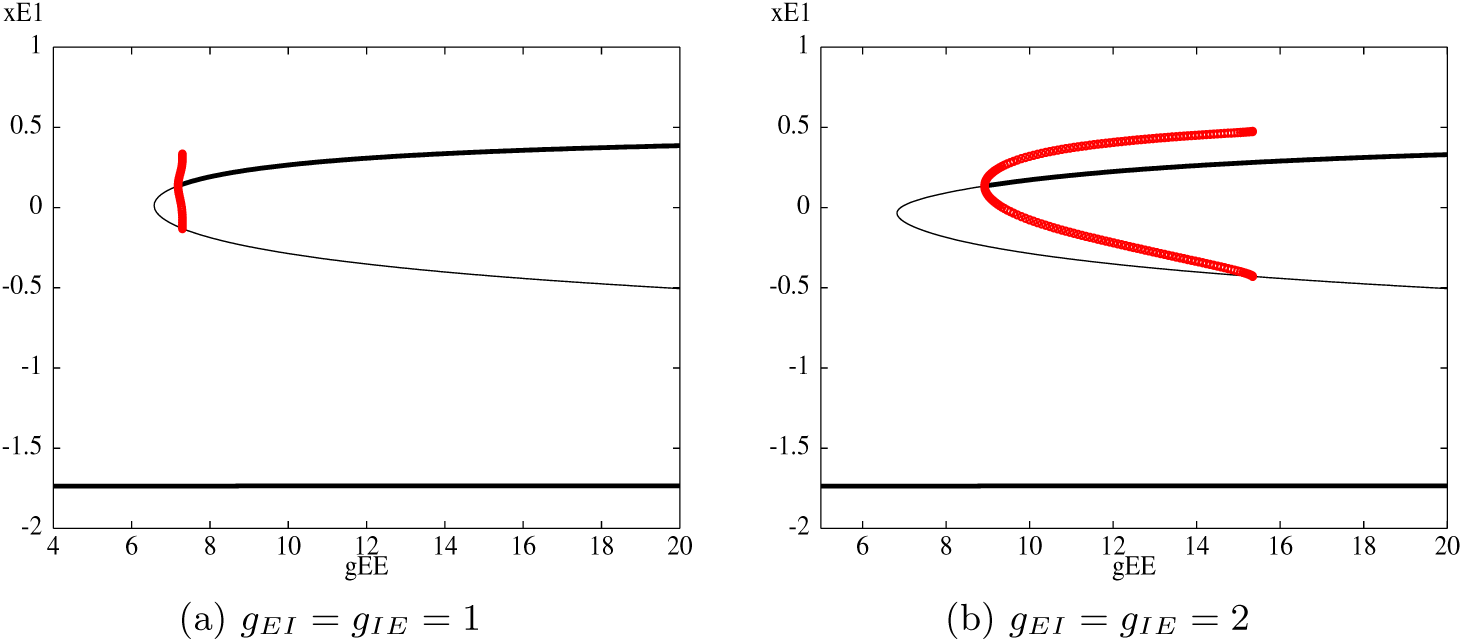
One parameter bifurcation diagrams *x*_*E*_ vs *g*_*EE*_ for the example system with *β* = 1.5, *I*_*E*_ = *I*_*I*_ = 0 and *g*_*EI*_, *g*_*IE*_ as shown. Other parameter values are given by (33). Thin/thick black curves show unstable/asymptotically stable equilibrium points. Red curves show maximum and minimum amplitude of unstable limit cycle.

**Figure 5:**
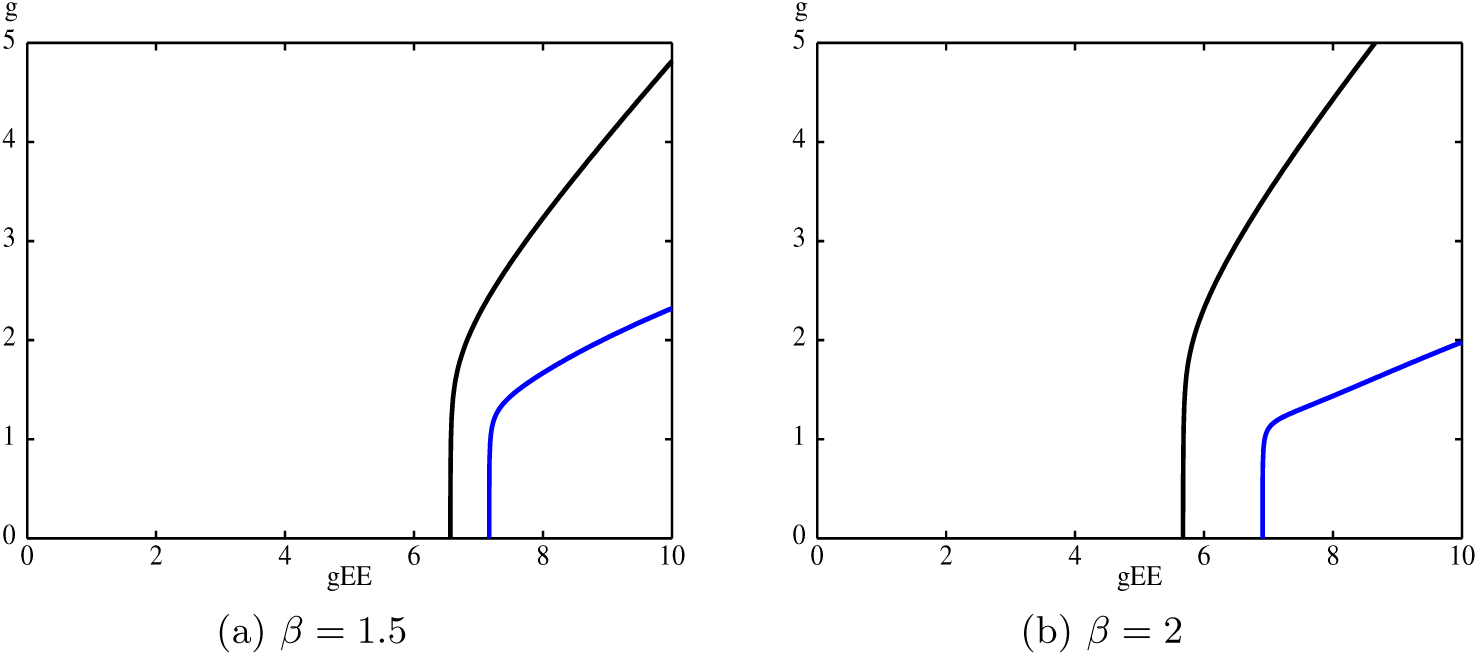
Two parameter bifurcation diagrams *g*(= *g*_*EI*_ = *g*_*IE*_) vs. *g*_*EE*_ for the example system. Parameter values are as given in (33) with *I*_*E*_ = *I*_*I*_ = 0 and with *β* as shown. Black is curve of saddle-node (fold) bifurcations. Blue is curve of Hopf bifurcations.

Next we considered the coupled system with delay, given by Eqs. (3)– (6), with the *F*_*E*_ = *F*_*I*_ = *F* and *G*_*E*_ = *G*_*I*_ = *G* given by Eq. (32) and the parameters as described above. Using Maple, we numerically solved for the equilibrium points as a function of *g*_*EE*_. This yielded the same curves of equilibrium points as in Fig. 4(a), which is to be expected since the equilibrium points do not depend on the size of the delay. Next we evaluated the Jacobian of the linearization at the equilibrium points and plotted the curves corresponding to Eqs. (26)–(27) as a function of *g*_*EE*_ for two different values of *g* = *g*_*IE*_ = *g*_*EI*_. These are shown in Fig. 6. The blue curves correspond to the highest equilibrium point in Fig. 4(a), with the thin curves corresponding to the Δ_+_ factor in (16) having a pair of pure imaginary eigenvalues and the thick to the Δ_*−*_ factor. The cyan curves correspond the middle equilibrium point. For all parameter values we considered, the middle equilibrium point was always unstable, thus these curves will not affect the observable dynamics. As noted above, for our choice of parameter values we expect the low (resting) equilibrium point to be asymptotically stable for *g*_*EE*_ in the range we consider. The black vertical line corresponds to the characteristic equation having a zero root. This value is independent of the delay and corresponds to the saddle-node bifurcation points in Fig. 4.

**Figure 6:**
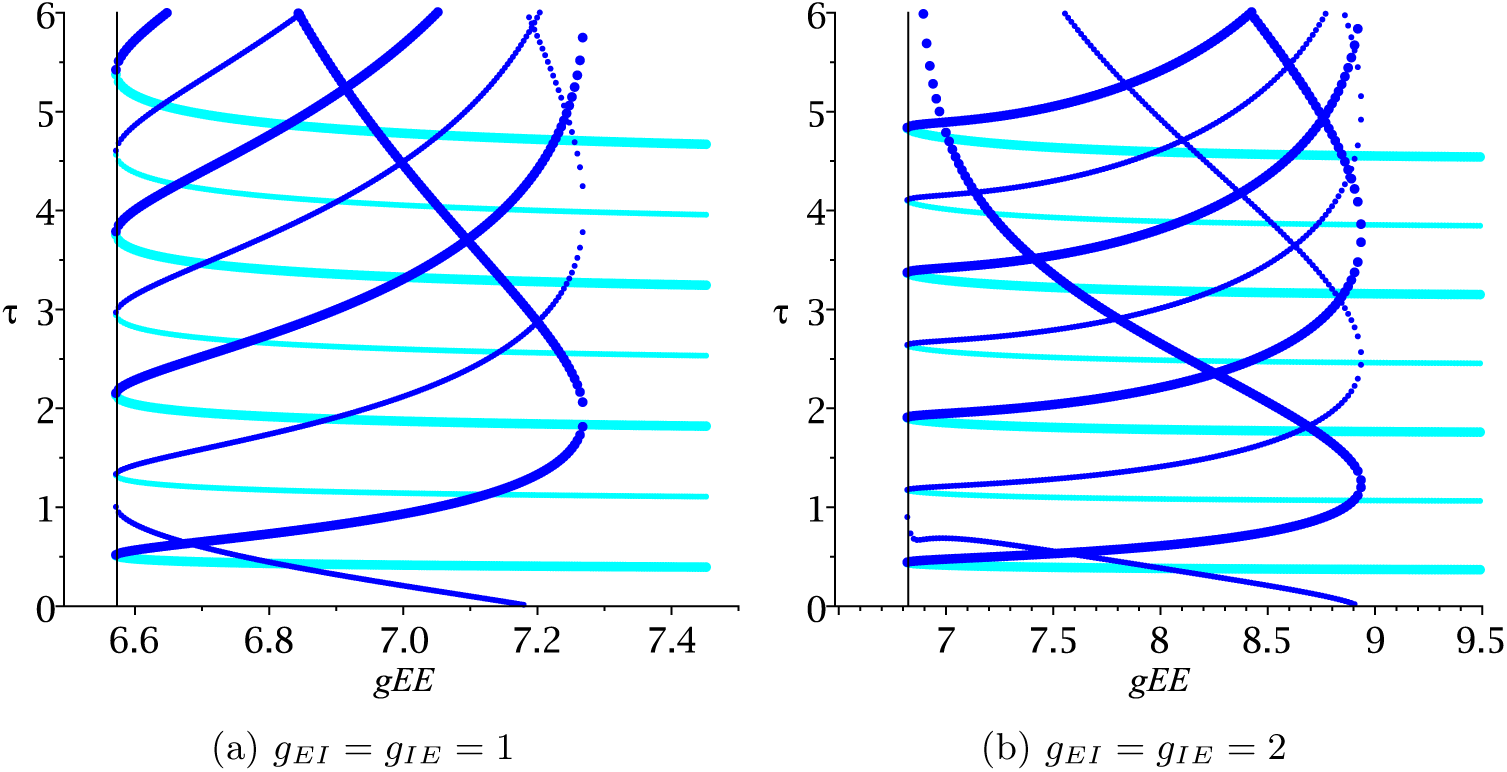
Plots of the curves in the *g*_*EE*_, *τ* parameter space where the characteristic equation (16) has a pair of pure imaginary eigenvalues. Blue curves correspond to the high equilibrium point, cyan to the middle equilibrium point. Thin curves correspond to Eq. (26) (roots of Δ_+_) thick to Eq. (27) (roots of Δ_*−*_). Parameter values are the same as Fig. 4(a), except *τ* > 0 and *β* is as given.

The high equilibrium point is asymptotically stable in the region to the right of all the blue curves. This follows from the fact that for *τ* = 0 the high equilibrium point is stable to the right of the intersection point of the lowest thin blue curve with the *g*_*EE*_ axis. This point corresponds to the Hopf bifurcation at 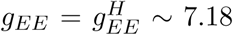 in Fig. 4(a) and at 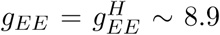 in Fig. 4(b). Consider Fig. 6(a). For any fixed 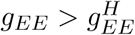 as *τ* is increased the high equilibrium point will lose stability at the first thick blue curve. If *g*_*EE*_ is large enough then no curve is crossed for any *τ* > 0 and the equilibrium point remains stable. If 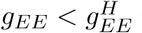 then the high equilibrium point will gain stability at the first thin blue curve then lose it at the first thick blue curve. The effect of inhibition can be seen by comparing Figs. 6(a) and (b). While qualitatively, the figures are similar, there are some quantitative differences. From a purely geometric perspective, larger *g*_*EI*_ and *g*_*IE*_ shift the Figure to the right, stretch it horizontally and compress it vertically. The biological implication is that if the *E*-*I* coupling is stronger then the stronger *E*-*E* coupling is needed to create the nonresting equilibrium points (as seen for zero delay in Fig. 4). However, smaller delays are needed to induce the Hopf bifurcations.

We expect that the high equilibrium point will undergo a Hopf bifurcation at *τ* values corresponding to the blue curves. The relative phases of the two cell populations on the resulting periodic orbits will be determined by the eigenvector structure. For example, if *τ*_1_ = *τ*_2_ then the eigenvector corresponding to a pair of pure imaginary roots of Δ_+_(*λ*) will be of the form (*ξ, ξ*)^*T*^. Thus the periodic orbits emanating from Hopf bifurcations along the thin blue curves will be of in-phase type: *x*_*E*1_(*t*) = *x*_*E*2_(*t*) and *x*_*I*1_(*t*) = *x*_*I*2_(*t*). Along the thick blue curves, the roots correspond to Δ_*−*_(*λ*) and the corresponding eigenvector is of the form (*ξ*, −*ξ*)^*T*^. Thus the corresponding periodic orbits will be of anti-phase type that is *x*_*E*1_(*t*) = *x*_*E*2_(*t* + *T/*2) and *x*_*I*1_(*t*) = *x*_*I*2_(*t* + *T/*2). This follows from symmetric Hopf bifurcation theory [29, 61, 59].

Finally, we note that the intersections of the blue lines correspond to points where the high equilibrium point has two pairs of pure imaginary eigenvalues, while the intersection points of the black line with the blue and cyan curves corresponds to points where there is an equilibrium point with one zero eigenvalue and a pair of pure imaginary eigenvalues. These correspond to Hopf-Hopf bifurcations and saddle-node(fold)/Hopf bifurcations, respectively [30, 40]. While such codimension two bifurcations are relatively rare in ordinary differential equation models, they are not unusual in delay differential equation models [3, 9, 10, 11, 59].

To verify the predictions about Hopf bifurcations and periodic solutions, we use the numerical continuation software DDE-BIFTOOL [20]. In addition to locating Hopf bifurcation points, this software can numerically compute the coefficient of the normal form to determine the criticality of the Hopf bifurcation. Applying the DDE-BIFTOOL to the model (3)–(6), with the functions in Eq. (32), the parameter values in (33) and *τ*_1_ = *τ*_2_, we confirmed the curves in Fig. 6 are indeed curves of Hopf bifurcation. The bifurcations of the high equilibrium point were found to be always subcritical while those of the middle equilibrium point were supercritical. The periodic orbits produced by the bifurcation of the middle equilibrium point are unstable since the equilibrium point is unstable before the bifurcation.

### 4.2. Periodic Solutions

We also used DDE-BIFTOOL to follow branches of periodic solutions emanating from the Hopf bifurcation points. Fig. 7(a) shows branches of periodic solutions aring from various branches of Hopf bifurcations at *g*_*EE*_ ≈ 7.2 using *τ*_*E*_ as the bifurcation parameter. Note that, the Hopf bifurcations are subcritical and produce unstable periodic orbits which coexist with the stable equilibrium point. However, in almost all cases, these branches undergo a saddle node of periodic orbits giving rise to stable periodic orbits. The far left curve in Fig. 7(a) starts from the Hopf point on lowest thick blue curve in Fig. 6(a) at *τ* ≈ 1.5. The far right curve in Fig. 6(a) (which remains unstable) starts from the same thick blue curve but at *τ* ≈ 2.7. The other curve of periodic solutions starts from the Hopf point on the thin blue curve at *τ* ≈ 3.2. Fig. 7(b) shows the form of the periodic solutions for *x*_*E*1_ and *x*_*E*2_ at particular points on the branches. Other points on the same branch produced different shapes of periodic orbits, but the phase relationship between *x*_*E*1_ and *x*_*E*2_ remains the same. Note that the phase relationships correspond with the predictions of our analysis, namely, the periodic solutions arising from the thick blue curves give rise to anti-phase periodic solutions and those arising from the thin blue curves to anti-phase solutions.

**Figure 7:**
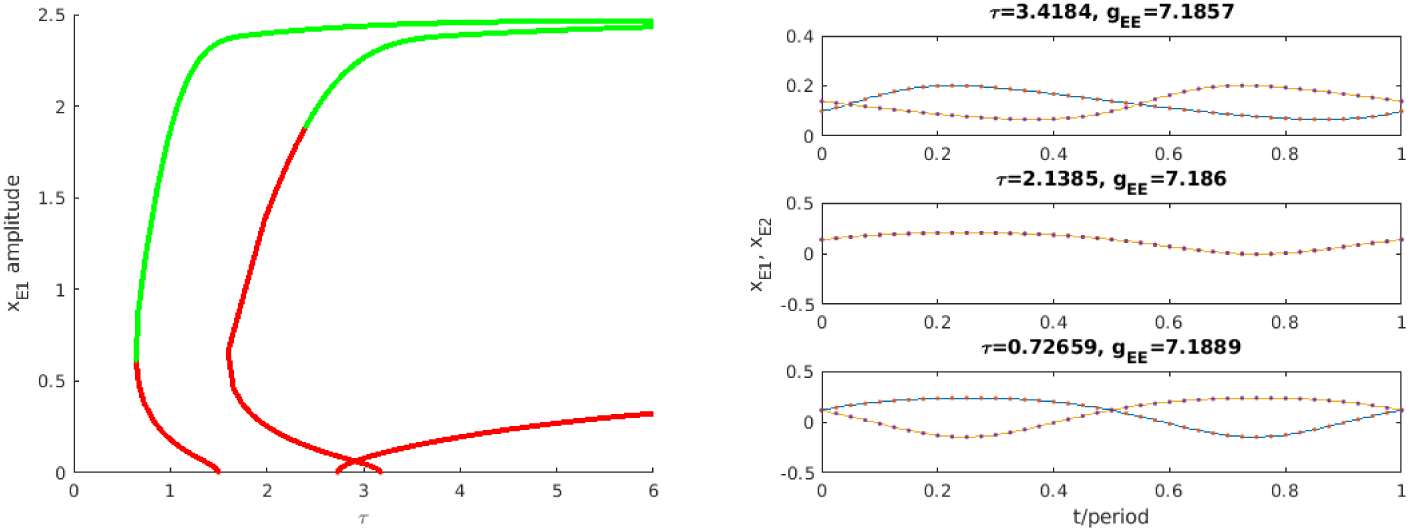
(a) Branches of periodic solutions with *g*_*EE*_ *≈* 7.2. Green/red points correspond to stable/unstable periodic solutions. (b) Periodic solutions for particular parameter values on each branch. Bottom corresponds to leftmost branch in (a); top to rightmost branch.

Figure 8 verifies this from a different perspective. It shows two branches of periodic solutions with *g*_*EE*_ as the bifurcation parameter, arising from to two different Hopf branches. The upper branch of periodic solutions in Fig. 8(a) corresponds to thin blue Hopf curve in Fig. 6(a) which passes through *τ* = 4.1864 when *g*_*EE*_ ≈ 7.23. The corresponding periodic solution in Fig. 8(b) is of in-phase type as predicted by our analysis. The lower branch of periodic solutions in Fig. 8(a) corresponds to the thick blue Hopf curve in Fig. 6(a) which passes through *τ* = 1.9184 when *g*_*EE*_ ≈ 7.23. The corresponding periodic solution in Fig. 8(b) is of anti-phase type as predicted.

**Figure 8:**
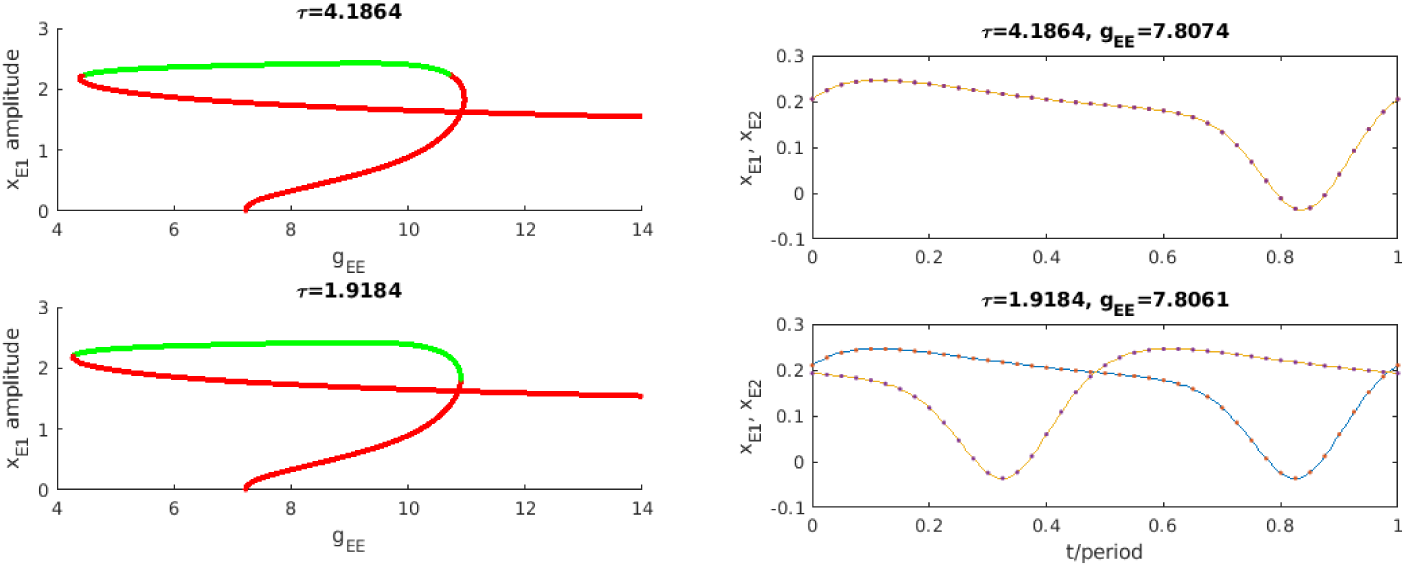
(a) Branches of periodic solutions with *g*_*EE*_ as the continuation parameter and *τ*_*E*_ as shown. Green /red points correspond to stable/unstable periodic solutions. (b) Periodic solutions for particular parameter values on each branch.

Finally, we confirmed the predictions of our analysis using numerical simulations of the model in XPPAUT [21]. Some examples are show in Fig. 9. Here we fixed 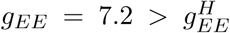. In all simulations the initial conditions were

**Figure 9:**
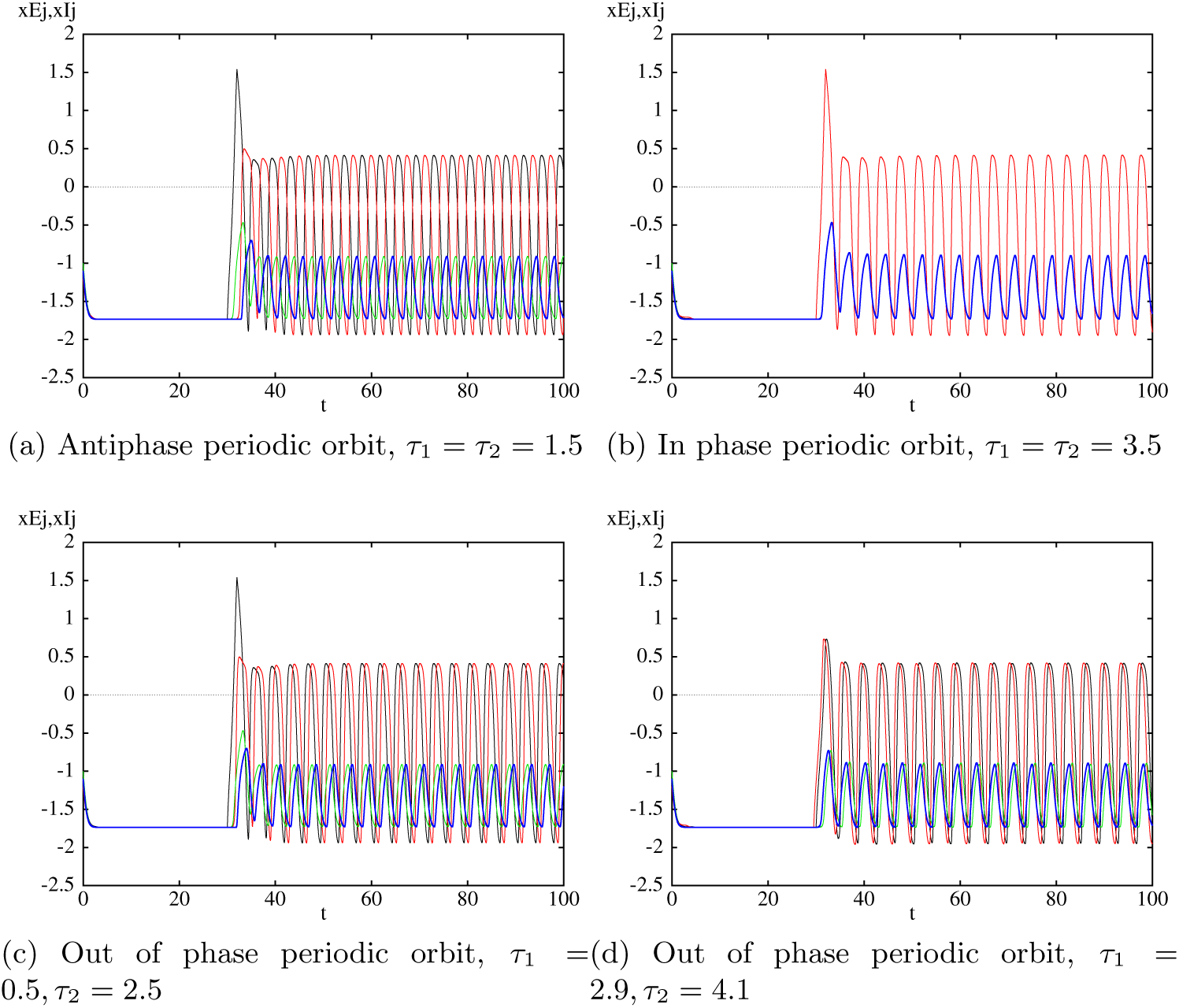
Numerical simulations of the model (3)–(6) with nonlinearities given by (32). *g*_*EE*_ = 7.2 and *τ*_1_, *τ*_2_ values as shown. Other parameter values are the same as Fig. 6(a). Black/red corresponds to *x*_*E*1_*/x*_*E*2_, while green/blue corresponds to *x*_*I*1_*/x*_*I*2_. Initial conditions are described in the text. Transient stimulation is applied as follows. (a) and (c) *E* cell 1: *I*_*app*_ = 2, 30 ≤ *t* ≤ 32; (b) both *E* cells: *I*_*app*_ = 2, 30 ≤ *t* ≤ 32; (d) *E* cell 1: *I*_*app*_ = 1.7, 30 ≤ *t* ≤ 32, *E* cell 2: *I*_*app*_ = 1.7, 29.5 ≤ *t* ≤ 31.5.

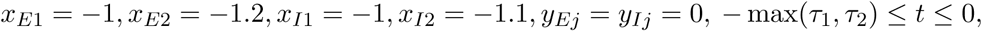

which leads to the low (resting) equilibrium point which is asymptotically stable. A brief stimulation, using the applied current *I*_*app*_ is applied (for details see Figure caption) and the solution switches to another asymptotically stable solution. For *τ* sufficiently small the high equilibrium point is asymptotically stable (not shown). At *τ* ∼ 1.3 (on the thick blue line) the high equilibrium point loses stability. If *τ*_1_ = *τ*_2_, the resulting periodic orbit is predicted to be of anti-phase type. Fig. 9(a) confirms this. As *τ* is increased further the thick blue line is crossed again (which restabilizes the equilibrium point) and then the think blue line is crossed. This latter results in a Hopf bifurcation leading to a periodic orbit which is predicted to be of in-phase type for the case *τ*_1_ = *τ*_2_. Fig. 9(b) confirms this. Note that both periodic orbits are large amplitude which is consistent Figs. 7–8. If *τ*_1_ ≠ *τ*_2_ but the Hopf bifurcations will still occur at the values of (*τ*_1_ + *τ*_2_)*/*2 indicated by Fig. 6, however, the phase relationship of the resulting periodic solution will be different. This is illustrated in Figs. 9(c) and (d) where solutions are shown which have with the same parameters and value of *τ*_1_ + *τ*_2_ as in Figs. 9(a) and (b), but *τ*_1_ ≠ *τ*_2_. In all cases, the solution can be switched from the oscillatory solution back to the resting equilibrium point by a transient stimulus to the inhibitory cells (not shown).

## 5. Discussion

In this paper, we study a model for a network of excitable neurons with excitatory and inhibitory synaptic coupling to investigate the role of coupling delays in the existence and stability of different types of oscillatory network behavior. Unlike many neural network models, a single, uncoupled neuron in our neural system is assumed to have zero applied current so that the synaptic coupling between neurons is the main factor to give rise to network behavior. We argue that the resting equilibrium point, i.e., the equilibrium where all the neurons are at their uncoupled rest state, should persist and remain asymptotically stable in the presence of coupling, but that sufficiently large excitatory coupling can give rise to new equilibria. We then use linear stability analysis of these equilibria to show that bifurcations are only expected when there is coupling between the excitatory neurons. We show that the stability only depends on the average of the delays between the excitatory neurons and give explicit expressions for values of the average delay where the equilibria lose stability via a pair of pure imaginary eigenvalues. We show that these expressions come in two types, which correspond to two factors in the characteristic equation. We then analyze the eigenvector structure to predict the type of oscillatory patterns that would emerge in the corresponding delay-induced Hopf bifurcations. To our knowledge, this is the first article to carry out stability and bifurcation analyses on such a system.

We apply our analytical results to an example model inspired by that of Terman and Wang [56]. We solve for the equilibria of the model and show that the resting point always persists, while for sufficiently large excitatory coupling a saddle node bifurcation occurs giving rise to two other equilibria. We find curves of potential Hopf bifurcation of these equilibria in the parameter space of the average delay and excitatory coupling strength, and show the two types described above alternate as the delay increases.

We supplement and verify our analytical results using numerical simulations and numerical bifurcation analysis. Using DDE-BIFTOOL [20] we verify that the curves of pure imaginary eigenvalues are indeed Hopf bifurcations. For the case of equal delays, the two types of Hopf bifurcation give rise to periodic solutions where corresponding cells are in-phase and anti-phase. We find sets of parameter values where both types of periodic solutions exist and are asymptotically stable. Using numerical simulations in XPPAUT [21], we show that if the delays are not equal, but the average delay remains the same, the oscillation patterns are transformed as predicted by the eigenvector analysis.

This study extends our previous work [51] in that the neural networks considered herein include two distinct pairs of excitatory and inhibitory neurons, which are coupled through excitatory synapses. Previously, we considered the global inhibitory network where an excitatory population is uncoupled but connected to a global inhibitory neuron only. Thus, our study exhibits other types of network behaviors such as anti-phase solutions in addition to synchronization among excitatory neurons. Here, we also consider a general network of non-relaxation oscillators unlike our previous study. Moreover, we extend our previous work by conducting numerical bifurcation analysis based on DDE-BIFTOOL, in which we show the coupling delay plays a significant role in generating various types of periodic solutions as a result of Hopf bifurcations.

An important finding in the current work is that the resting equilibrium point is unaffected by the coupling. This is due to the fact that the synaptic coupling is effectively zero when the neuron is at rest, which is a common feature of biophysical models of neurons. Thus all the delay-induced bifurcations involve nontrivial equilibria which are induced by the excitatory coupling and the resultant stable periodic solutions coexist with the stable resting equilibrium. This is in contrast to studies of neural networks with delayed diffusive or sigmoidal coupling where the resting/trivial equilibrium point undergoes the delay-induced bifurcations and becomes unstable [8, 9, 10, 11, 15, 17, 18, 52, 41, 59].

Our results about Hopf bifurcation leading to in-phase and anti-phase periodic solutions, for the case of symmetric delays in the *E*-*E* connections, mirror those found in many systems of two coupled identical oscillators [9, 10, 15, 17, 41, 44, 52, 54]. Here we have shown that these results persist in models with synaptic coupling and even if the resting (trivial) equilibrium remains asymptotically stable. Further, we have shown how these results may be extended to the case of non-symmetric delays.

Our results have an interesting implications for how rhythms may arise in biological neural networks. Rhythms are associated with frequencies that are observed at the network level as opposed to the individual neuron level. One way that these can arise is through phase-locked oscillatory solutions [35, 37]. For example, in our small network in-phase oscillations give rise to a network frequency equal to the frequency of the individual neurons, while anti-phase oscillations give rise to a network frequency twice that of the individual neurons. Several studies have shown that rhythms can arise in networks of oscillatory neurons [35, 37], however, these rhythms are generally always present unless a parameter changes. In our model, due to the bistability between the resting equilibrium and the oscillatory solutions, the network can transition between a non-rhythmic state and a rhythmic state, simply through a transient input. Further, we observe parameter ranges where there is tristability between the resting equilibrium point and two different oscillatory patterns (in-phase and anti-phase). In this parameter regime the system can transition from the non-rhythmic activity to two different rhythmic states, depending on the input.

## Model limitations and future directions

Despite the richness of our analytical and numerical results, our study is based on the small-size network with four, two excitatory and two inhibitory, neurons only resulting in two excitatory-inhibitory pairs. The results for two neurons with symmetric delays have been extended to larger networks with circulant coupling in [59]. We expect that we can use a similar approach to extend our results for non-symmetric delays to these types of larger networks.

In addition, there is only one location where coupling delays are assumed to arise, which is in synapses between excitatory cells in different populations. Though our study considers asymmetric delay case among excitatory neurons but additional delays between coupled excitatory and inhibitory cells are needed to obtain a more complete understanding of a realistic neural network. We could also include delays within a population to better represent the synaptic delay between neurons. Further, in some brain networks there exist long-range connections from excitatory cells in one population to inhibitory cells in the other [34, 37]. If such connections were added to our model, it would be necessary to include the corresponding coupling delay. Thus, our future study would extend the current model to investigate the impact of delays between different types of neurons on the observed network behaviors in this study.

Our work here focused on models with one “activity” variable and one “gating” variable. However, as shown in [59] the linear stability analysis we use could be applied to more general conductance-based models with multiple gating variables.

Finally, real neurons are not likely to be perfectly symmetric. In other studies without delay, it has been shown that some results for symmetric systems persist, but new phenomena can also occur [12, 53]. Thus it would be interesting to include the effects of heterogeneity in neural parameters in a future study.

## Acknowledgments

SAC acknowledges the support of the Natural Sciences and Engineering Research Council of Canada.

## 6. Appendix Single Cell Model

In this paper we consider a single cell model inspired by that of Terman and Wang [56]. The model is based on the FitzHugh-Nagumo [24, 42] model but with a nonlinearity in equation for the “recovery” variable which is similar to that for a gating variable in a conductance-based model. The equations for this model are as follows

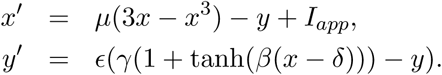

This nonlinearity for the recovery variable means that the model may act either as a class I and class II oscillator, depending on the parameter values.

For the parameters used in this study, that is, given in (33), the parameter *β* can be used to switch the model from a class 2 to a class 1 oscillator, with *I*_*app*_ used as the bifurcation parameter. This can be seen in Figure 10, which shows a one parameter bifurcation diagram of the system when *I*_*app*_ is used as the bifurcation parameter.

**Figure 10:**
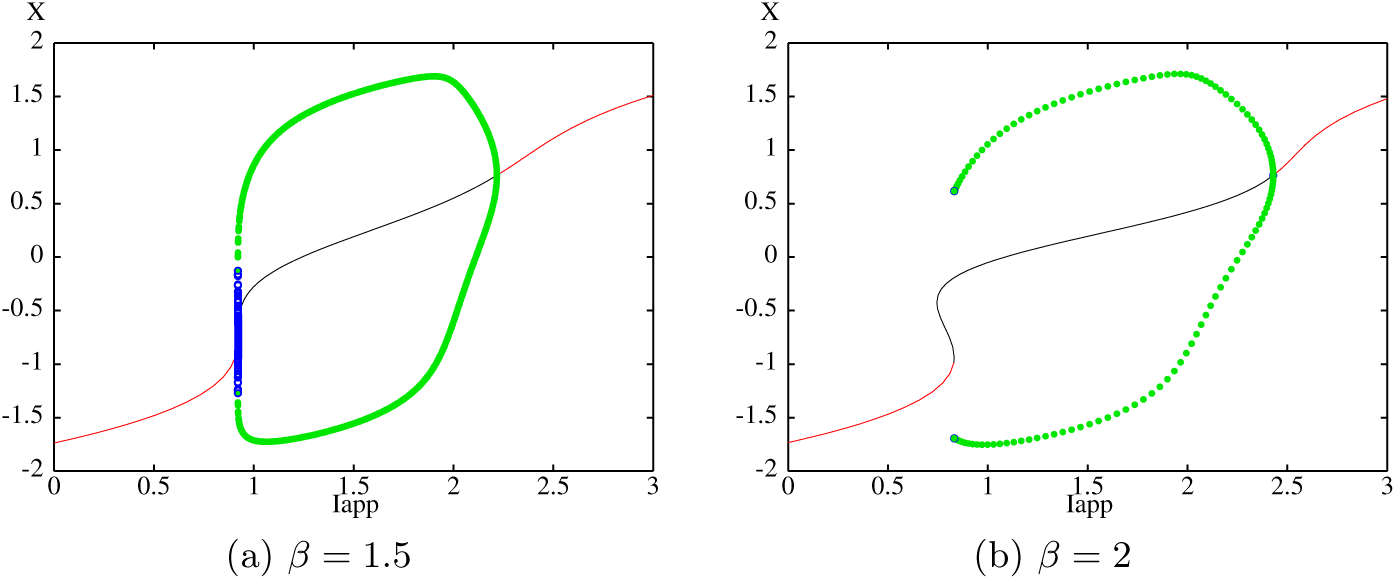
One parameter bifurcation diagram *V* vs. *I*_*app*_ with parameters given by (33) and *β* values as shown. Black/red curves correspond to asymptotically stable/unstable equilibrium points. Green/blue circles correspond to asymptotically stable/unstable periodic orbits.

When *β* = 1.5 there is a unique equilibrium point which goes loses stability in a subcritical Hopf bifurcation. A stable periodic orbit is created by a saddle node of periodic orbits. When *β* = 2.0 there are up to three equilibrium points and a stable periodic orbit is created in a saddle node on an invariant circle bifurcation. In both cases the cell is excitable for *I*_*app*_ small enough, exhibits stable spiking behavior (has a stable limit cycle) for a mid range of *I*_*app*_ values and has a “high amplitude” stable equilibrium point for *I*_*app*_ large enough. Typical spiking solutions are shown in Figure 11. Note that the *x* varies in the range [−2, 1] for these solutions.

**Figure 11:**
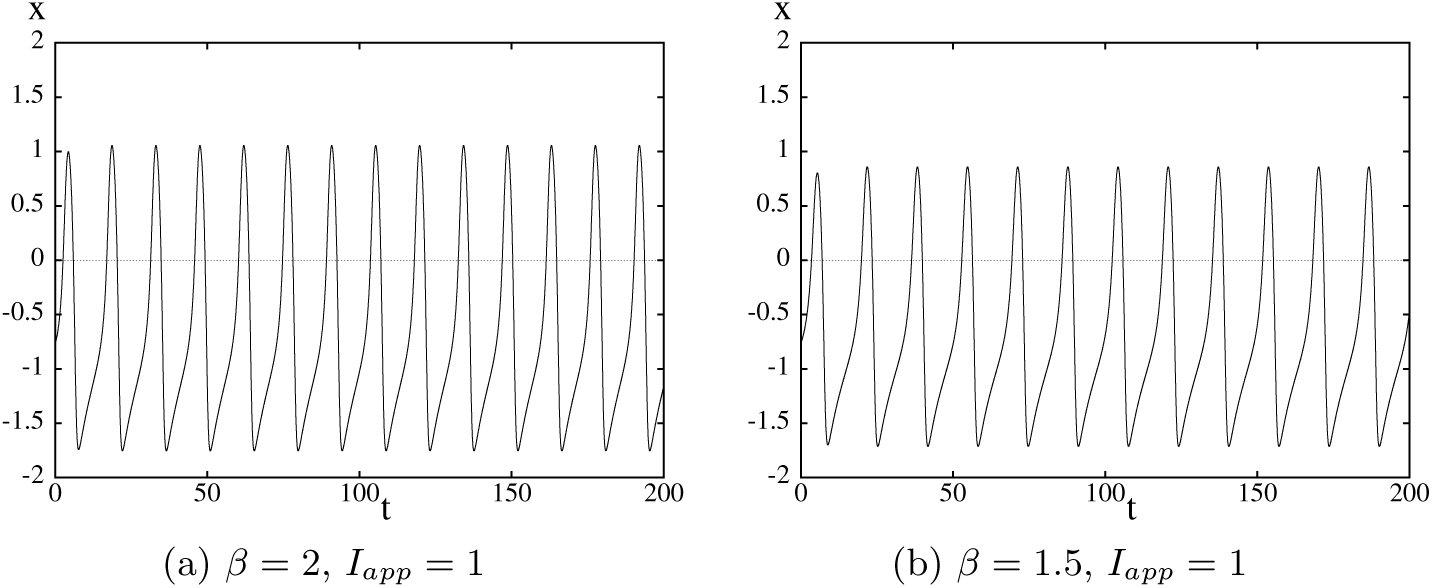
Typical spiking solutions of single cell model. Parameter values are given by (33) and *β* and *I*_*app*_ as shown.

